# Pinaceae show elevated rates of gene duplication and gene loss that are robust to incomplete gene annotation

**DOI:** 10.1101/192203

**Authors:** Claudio Casola, Tomasz E. Koralewski

**Author notes:** Corresponding author: 495 Horticulture Rd, College Station, 77843-2138 TX.

## Abstract

Gene duplications and gene losses are major determinants of genome evolution and phenotypic diversity. The frequency of gene turnover (gene gains and gene losses combined) is known to vary between organisms. Comparative genomic analyses of gene families can highlight such variation; however, estimates of gene turnover rates may be biased when using highly fragmented genome assemblies resulting in poor gene annotations. Here, we address potential biases introduced by gene annotation errors in estimates of gene turnover frequencies in a dataset including both well-annotated angiosperm genomes and the incomplete gene sets of four Pinaceae including two pine species, Norway spruce and Douglas-fir. Previous studies have shown low overall substitution rates, but higher levels of adaptive substitutions, in genes from Pinaceae and other gymnosperms compared to angiosperms. Conversely, our analysis suggests that pines experienced higher gene turnover rates than angiosperm lineages lacking recent whole-genome duplications. This finding is robust to both known major issues in Pinaceae gene sets: missing gene models and erroneous annotation of pseudogenes. A separate analysis limited to the four Pinaceae gene sets confirmed an accelerated gene turnover rate in pines compared to Norway spruce and Douglas-fir. Our results indicate that gene turnover significantly contributes to genome variation and possibly to adaptation and speciation in Pinaceae. Moreover, these findings indicate that reliable estimates of gene turnover frequencies can be discerned in incomplete and potentially inaccurate gene sets.

## Introduction

The processes of gene duplication and gene loss are major evolutionary events with profound effects on the emergence of novel phenotypes and organismal adaptation (Demuth and Hahn 2009; Kondrashov 2012; Albalat and Canestro 2016; Van de Peer, et al. 2017). The analysis of genome-wide datasets in a phylogenetic framework is a powerful tool to investigate gene duplication and gene loss events (i.e., gene turnover) that contributed to the evolution of lineage-specific traits. For instance, this type of studies has shown that gene turnover rates are overall comparable across mammals, *Drosophila*, yeasts and plants (Lynch and Conery 2000; Gu, et al. 2002; Hahn, et al. 2005; Maere, et al. 2005; Hahn, Han, et al. 2007; Guo 2013; Han, et al. 2013). At a finer evolutionary scale, however, some intriguing differences have emerged, such as a higher rate of gene duplication and loss in the lineage leading to modern humans compared to other branches of the great apes phylogeny (Hahn, Demuth, et al. 2007; Marques-Bonet, et al. 2009; Locke, et al. 2011; Carbone, et al. 2014). Therefore, the pace of gene duplication and gene loss may differ significantly even between closely related species, arguably fueling phenotypic divergence and adaptation.

The availability of draft genome sequences from a rapidly increasing variety of organisms offers the unprecedented opportunity to investigate at multiple taxonomic scales the tempo and mode of gene turnover. In spite of their potential to provide crucial information on the emergence of phenotypic innovation, analyses of gene turnover can be severely biased because of the presence of incomplete and/or potentially inaccurate gene datasets. For instance, erroneous gene losses will be observed whenever genes occurring in a genome are not annotated in the corresponding gene set, whereas apparent gene duplications will be reported whenever single genes are broken down into multiple gene models in highly fragmented genome assemblies, or whenever the two alleles of a gene are annotated as two separate loci (Denton, et al. 2014). Although several computational approaches have been developed to provide estimates of gene annotation errors (Parra, et al. 2007; Denton, et al. 2014; Simao, et al. 2015), and a few bioinformatic tools currently available can overcome some limitations in gene annotations (Zhang, et al. 2016), the correct prediction of the gene space remains one of the most problematic outcomes of modern eukaryotic genomics, particularly but not exclusively in non-model organisms. This issue is especially acute in taxa with very large genomes, which tend to be represented by assemblies with a high number of short contigs and scaffolds (Florea, et al. 2011; Denton, et al. 2014).

The family Pinaceae, with their haploid genome sizes ranging between ∼10 and ∼36 Gb (De La Torre, et al. 2014), is one of such groups. Owing to a combination of sequencing techniques and, more importantly, to a series of novel bioinformatic strategies and tools, the enormous genomes of several Pinaceae have been recently sequenced and assembled (Birol, et al. 2013; Nystedt, et al. 2013; Neale, et al. 2014; Wegrzyn, et al. 2014; Zimin, et al. 2014; Warren, et al. 2015; Gonzalez-Ibeas, et al. 2016; Stevens, et al. 2016; Neale, et al. 2017). However, given the size and repetitiveness of these genomes, the resulting assemblies tend to be highly fragmented, with a negative effect on the accuracy of gene annotation.

Additionally, Pinaceae genomes contain a high number of ‘gene fragments’— short pseudogenes derived from duplication and retention of one or a few exons of functional genes (Nystedt, et al. 2013; Wegrzyn, et al. 2014). These gene fragments add another layer of complexity in the gene annotation process. Many gene fragments appear to share a high sequence identity with their parental genes and show no disabling mutations; therefore, such short pseudogenes may form a substantial portion of the annotated gene models in Pinaceae.

In this study, we generated gene family datasets to estimate and compare rates of gene turnover across four Pinaceae, six dicots, five monocots and the moss *Physcomitrella patens*, using a maximum-likelihood framework implemented in the package CAFE (Hahn, et al. 2005). This approach allowed inferring the size of each gene family at ancestral nodes of the analyzed phylogeny (Han, et al. 2013). For each branch of the tree, gene duplications and gene losses were then calculated by comparing the size of gene families at the two nodes delimiting that branch. This approach is particularly suitable to identify small-scale gene duplications and losses, as opposed to gene turnover events associated with polyploidization, or whole-genome duplications (WGD). This is a consequence of the widespread and relatively rapid removal of one of the two homeologous gene copies following WGD (Freeling 2009). Therefore, gene turnover rates might be underestimated using such method, particularly in lineages that underwent multiple rounds of WGD.

Gymnosperms and angiosperms underwent a polyploidization event before their separation (Jiao, et al. 2011), but the number and timing of subsequent WGDs vary substantially across modern seed plant groups. In Pinaceae, a single ancient WGD appears to have occurred between 210 and 342 million years ago (Li, et al. 2015). Conversely, many angiosperm lineages have experienced one or multiple recent WGDs, defined here as polyploidization events that occurred in the past 75 million years (Jiao, et al. 2011; Vanneste, et al. 2014; Soltis, et al. 2015; Wendel 2015). However, flowering plant lineages such as those leading to modern cacao, peach and castor bean, three species included in this study, did not go through polyploidization after the γ WGD event, estimated to have occurred around 120 million years ago (Chan, et al. 2010; Argout, et al. 2011; Jiao, et al. 2011; International Peach Genome, et al. 2013; Li, et al. 2016). Therefore, modern Pinaceae and the dicots cacao, peach and castor bean share the same number of WGD events.

Testing evolutionary models that incorporate separate estimates of gene turnover rates in Pinaceae, angiosperms with recent WGDs and the three dicots without recent WGDs revealed that Pinaceae experienced rates of gene duplication and gene loss comparable to those of flowering plants. Correcting for incomplete gene annotation and erroneous pseudogenes annotation in the Pinaceae gene sets did not alter significantly these results. Among Pinaceae, we found a higher frequency of gene gains and losses in two pines, *Pinus taeda* (loblolly pine) and *Pinus lambertiana* (sugar pine) compared to Norway spruce (*Picea abies*) and Douglas-fir (*Pseudotsuga menziesii*). These findings are in contrast to the reported very low nucleotide substitution rates in Pinaceae (Willyard, et al. 2007; Buschiazzo, et al. 2012; Chen, et al. 2012; De La Torre, et al. 2017) and suggest that the frequency of gene duplication and gene loss events is uncoupled from the per-base substitution rate in Pinaceae. Thus, gene turnover represents a fundamental source of genome novelty and potentially adaptation in pines, spruces and Douglas-fir.

## Results

### Gene family datasets

Two primary gene family datasets were generated. First, unique protein sets of the 16 analyzed plant species (Fig. 1) were used to obtain orthogroups, i.e. gene families, with OrthoFinder (Emms and Kelly 2015). This dataset included 66-96% of all the original genes depending on the species (Table 1). Compared to other land plants analyzed, Pinaceae showed a higher number of total genes and genes clustered in gene families. A subsequent filtering step aiming at removing gene families that are largely lineage specific (see Methods) resulted in a final dataset that had a comparable number of genes in 8,698 gene families across the 16 species (Table 1). We named this group the land plant gene family (LGF) dataset.

**Figure 1.**
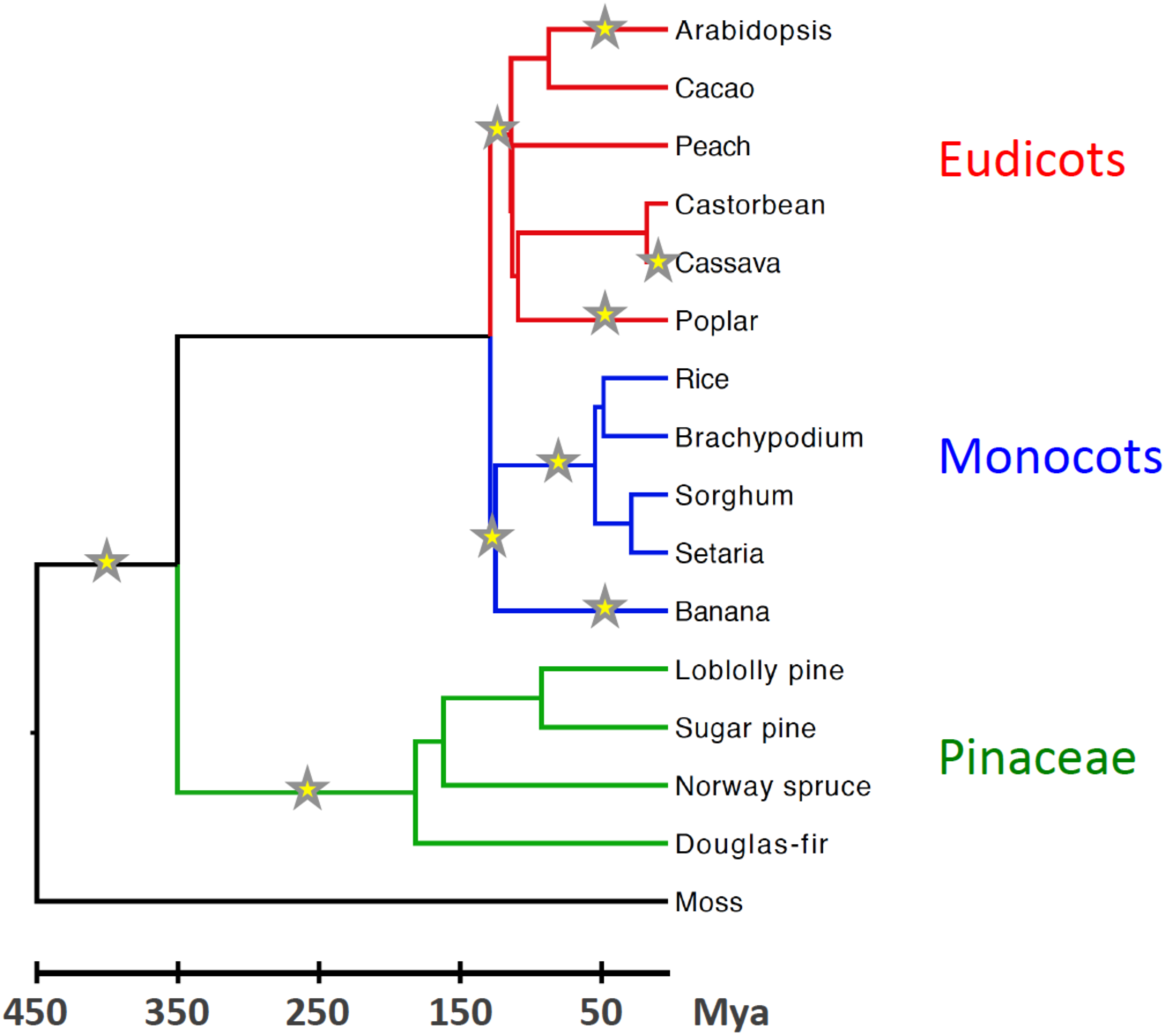
Phylogeny of the sixteen land plants analyzed in this study. Background branches are in black, eudicot branches – red, monocot– blue, and Pinaceae – green. Branches that experienced whole-genome duplication (WGD) events are highlighted by a star. Multiple WGDs on a single branche are not shown.

**Table 1.** Analyzed species and genes in the land plant datasets.

| Name | Species name | Number of genes | Number of genes in orthogroup<sup>a</sup> | Number of genes in LGF<sup>a</sup> | Number of genes in LGF50<sup>a</sup> |
| --- | --- | --- | --- | --- | --- |
| Arabidopsis | *Arabidopsis thaliana* | 26930 | 23711 (88.1) | 16551 (61.5) | 16216 (60.2) |
| Cacao | *Theobroma cacao* | 29400 | 24201 (82.4) | 14639 (49.8) | 14166 (48.2) |
| Peach | *Prunus persica* | 27861 | 25525 (91.7) | 15004 (53.9) | 14473 (51.9) |
| Castor bean | *Ricinus communis* | 31167 | 21800 (70.0) | 14222 (45.7) | 13584 (43.6) |
| Cassava | *Manihot esculenta* | 30666 | 29319 (95.7) | 19767 (64.5) | 18752 (61.1) |
| Poplar | *Populus trichocarpa* | 41234 | 35340 (85.8) | 22769 (55.3) | 21614 (52.4) |
| Rice | *Oryza sativa* | 39004 | 27865 (71.5) | 16047 (41.2) | 15533 (39.8) |
| Brachypodium | *Brachypodium distachyon* | 31560 | 27087 (85.9) | 15900 (50.4) | 15497 (49.1) |
| Sorghum | *Sorghum bicolor* | 32915 | 27325 (83.1) | 16511 (50.2) | 16073 (48.8) |
| Setaria | *Setaria italica* | 33543 | 28908 (86.2) | 16547 (49.4) | 15935 (47.5) |
| Banana | *Musa acuminata* | 34329 | 28777 (83.9) | 22014 (64.2) | 20394 (59.4) |
| Loblolly pine | *Pinus taeda* | 83861 | 57768 (68.9) | 18342 (21.9) | 13525 (16.1) |
| Sugar pine | *Pinus lambertiana* | 84988 | 62175 (73.2) | 17782 (21.0) | 13637 (16.0) |
| Norway spruce | *Picea abies* | 63621 | 47904 (75.3) | 20649 (32.5) | 15616 (24.5) |
| Douglas-fir | *Pseudotsuga menziesii* | 54626 | 43382 (79.5) | 15524 (28.5) | 12788 (23.4) |
| Moss | *Physcomitrella patens* | 26491 | 17465 (66.0) | 14016 (53.0) | 13460 (50.8) |
| **Total** |  | 672196 | 528552 | 276284 | 251263 |

<sup>a</sup> Percentages are shown in parenthesis

A Pinaceae gene family (PGF) dataset was also generated using only genes from the four Pinaceae species (Table S1). Clustering with OrthoFinder and subsequent filtering out the lineage-specific orthogroups resulted in 16,118 gene families, almost twice as many as in the LGF dataset, despite the lower number of Pinaceae genes. This is likely due to a different clustering of genes in datasets with four versus sixteen species. Indeed, the average number of genes per gene family in the four Pinaceae shifts from 1.6 in the PGF to 2.1 in the LGF (data not shown).

Pinaceae genomes have been shown to contain thousands of short pseudogenes that may be included in gene sets (Nystedt, et al. 2013; Wegrzyn, et al. 2014). Additionally, the fragmentation of genome assemblies in these conifers leads to many genes being broken into multiple gene models, a phenomenon that has been referred to as gene cleavage (Denton et al. 2014). These are reasons that likely contribute to the observed decreased length of Pinaceae proteins as compared to other LGF species (Fig. S1A).

To minimize potential bias in gene duplication rates due to erroneous pseudogene annotation and gene cleavage in Pinaceae we analyzed two smaller versions of the LGF and PGF datasets wherein proteins with a sequence length ≤50% of the longest protein in the same gene family from the same species were removed. These two versions are referred to as LGF50 and PGF50 gene sets. In angiosperms, the LGF50 gene set differs only slightly from the LGF set (∼2-7% of genes were removed), whereas in Pinaceae 18-26% of genes were parsed out from the LGF set (Table 1; Fig. S2). As a result, the gap in the protein length distribution between Pinaceae and other plants is markedly reduced in the LGF50 (Fig. S1B). Comparably large contractions of gene families were found between the PGF and the PGF50 gene sets (Table S1).

### Pinaceae have higher gene turnover rates than angiosperm lineages with no recent whole-genome duplications

To infer patterns of gene family evolution in Pinaceae we applied the maximum-likelihood framework implemented in the CAFE package (Han, et al. 2013) on the four datasets described above. In CAFE, the parameter λ represent the gene turnover rate (λ = number of gene gains and losses/gene/million years). Multiple λ values can be estimated on branches of the phylogenetic tree of interest in order to test hypothesis on the variation of gene turnover rates between taxa.

We first compared overall gene turnover rates in a four-λ model with separate rate estimates for dicots, monocots, Pinaceae and a group of background branches also including the moss lineage (Fig. 2; Table S2). In this model, Pinaceae gene turnover rates are 1.31, 1.56 and 5.40 fold higher than dicots, monocots and background branches, respectively, when using the LGF dataset (Fig. 2A; Table S2). The Pinaceae frequency of gene turnover decreased to 0.96/1.12/3.56 times that of the three other lineages in the same model incorporating the LGF50 dataset (Fig. 2B; Table S2).

**Figure 2.**
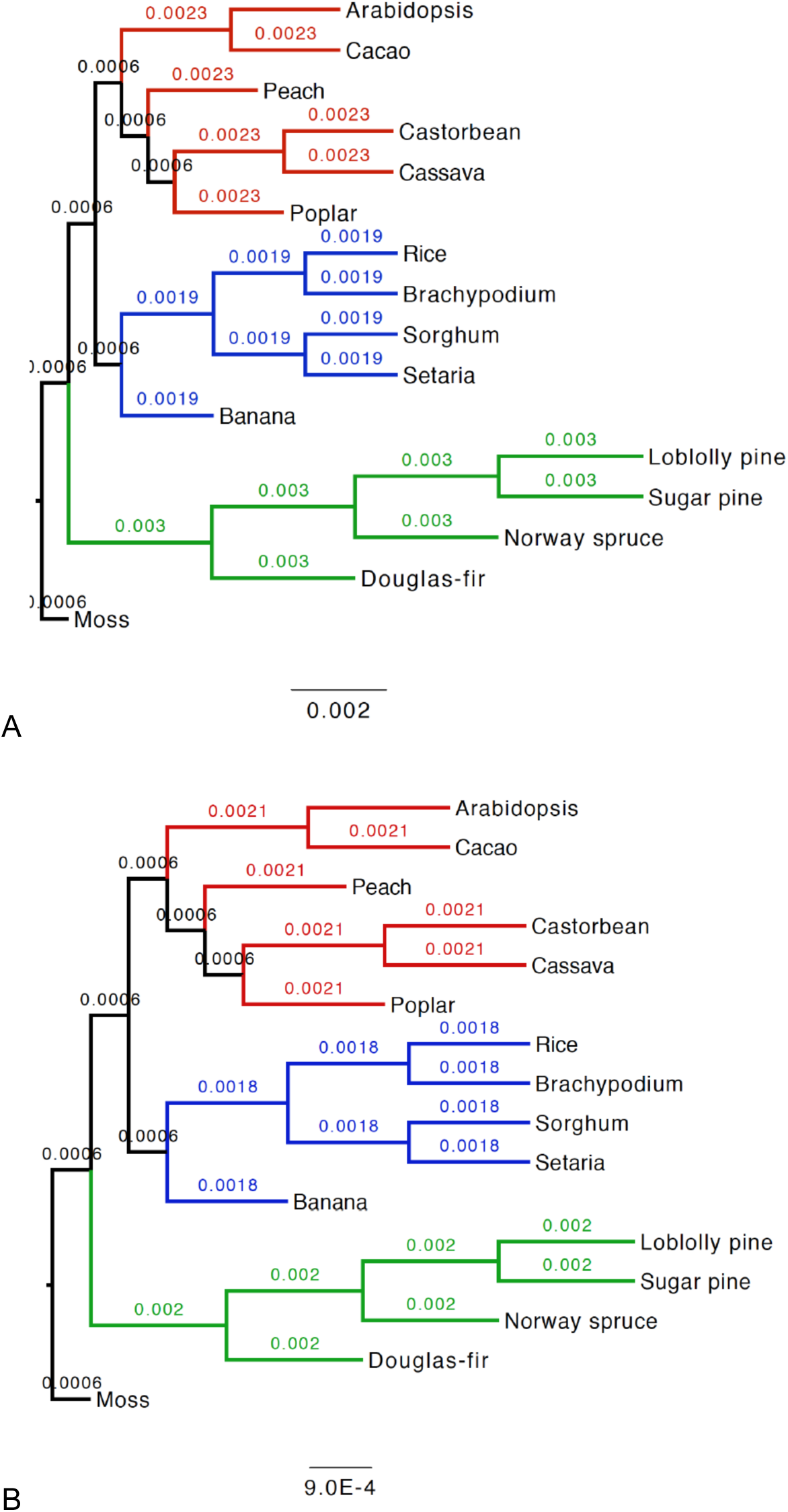
Gene turnover rates in four-λ models applied to the LGF (A) and LGF50 (B) gene sets. Dicots, monocots and Pinaceae branches are color-coded as in Figure 1.

To further assess gene gain and loss frequencies across taxa we examined phylogenies where 5 to 9 λ values were calculated. As expected, the likelihood values of the models implemented in the CAFE analysis tended to increase with more λ values (data not shown). However, convergence in models with more than six λ values was achieved only in a few cases, possibly because of overparameterization. Therefore, we carried out an extensive analysis of fourteen models estimating six λ values aimed at determining gene turnover rates in lineages of Pinaceae and dicots with and without recent WGD (Table S3). These fourteen models were chosen among all possible six-λ models in order to obtain separate gene turnover estimates for each major taxonomic group, namely pines, Pinaceae, all dicots, dicots with recent WGDs, dicots without recent WGDs and all monocots, while grouping ancestral branches of the phylogeny together in a single λ category.

Minimum and maximum gene turnover rates were inferred for each branch in the phylogeny with both LGF and LGF50 gene sets. Pines exhibited both the highest minimum and maximum rates (Fig. 3; Tables S4-5). High rates were also observed in the ancestral pine branch, which result from models wherein this branch’s estimated λ value was the same of the two pine species λ.

**Figure 3.**
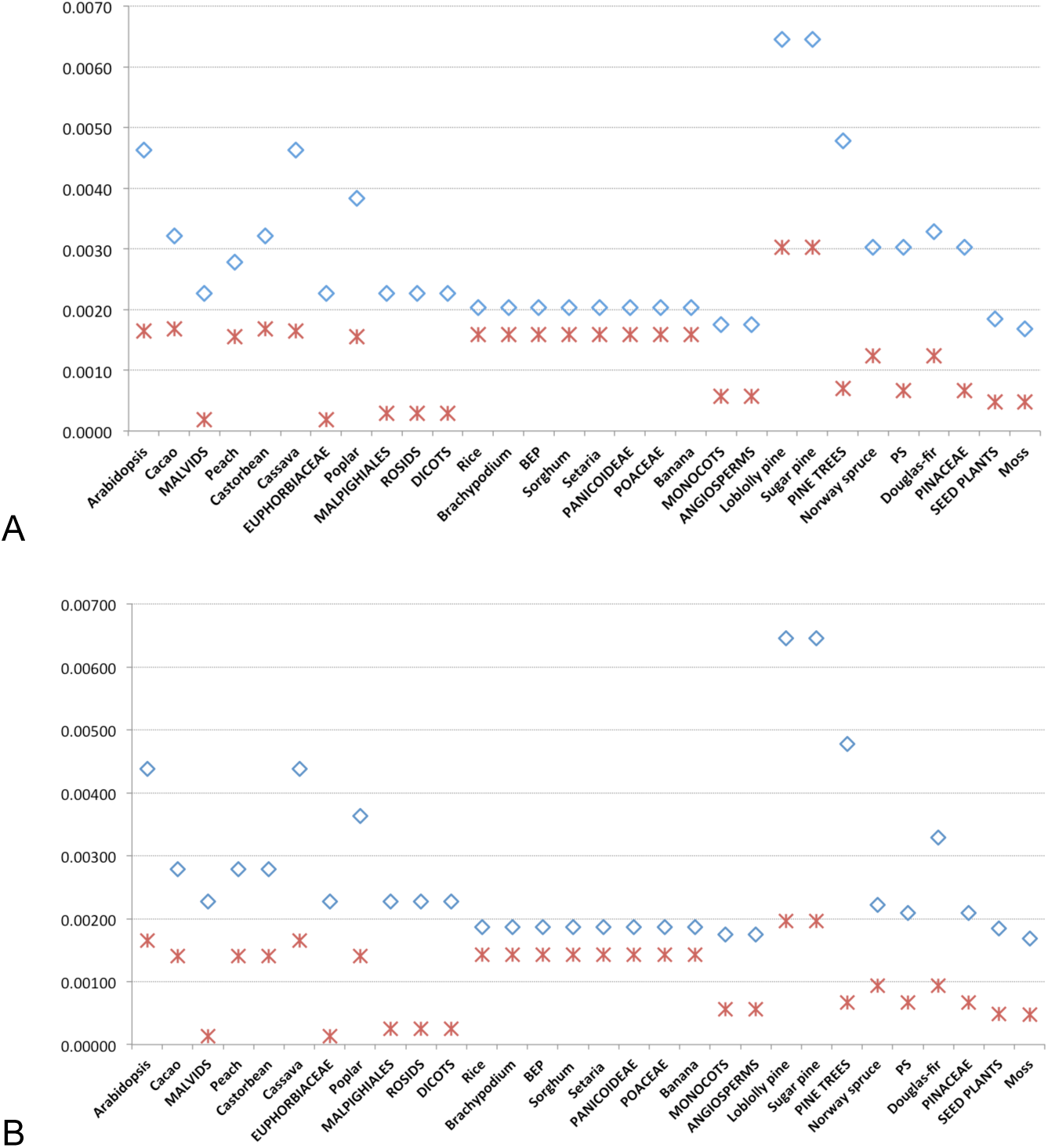
Maximum (diamonds) and minimum (asterisks) gene turnover rates in fourteen six-λ models applied to the LGF (A) and LGF50 (B) gene sets.

**Figure 4.**
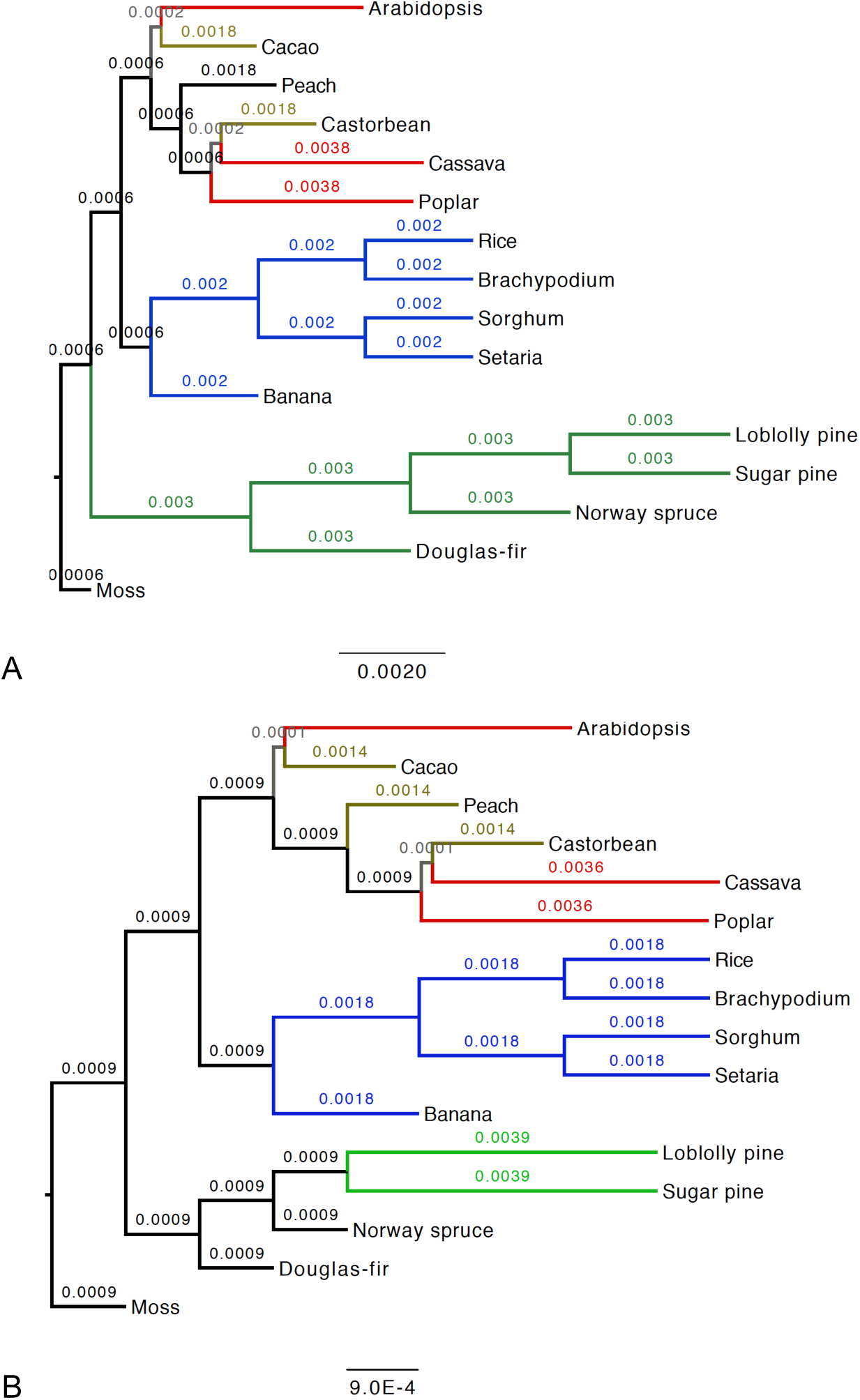
Gene turnover rates in the best fitting six-λ models applied to the LGF (A) and LGF50 (B) gene sets. Dicots with recent WGDs (red), dicots without recent WGDs (light brown), monocots (blue), all Pinaceae (dark green), pines (light green), dicots inner branches (grey) and background branches (black) are shown.

Among angiosperms, the three species that experienced recent WGDs—*Arabidopsis*, cassava and poplar—showed higher maximum gene turnover rates than the ancient-only WGD species cacao, castor bean and peach (Fig. 3). These results suggest that polyploidization leads to an increase in detected gene turnover events even when gene duplications and gene losses due to WGD are largely unidentified due to the node-to-node gene family size estimates. The retention of a sizable number of gene duplicates between ancestral and recent nodes of branches with WGD is probably responsible for the high levels of gene turnover in these species.

The total number of gene gains and losses was calculated for each of the sixteen species in the phylogeny. In the two pine species, 54-118 gene duplication/loss events occurred according to the minimum and maximum estimated gene turnover rates, respectively, in the LGF gene set (Fig. S3A; Table S4). The third highest number of gene changes was found in cassava with 33-92. Qualitatively similar results were observed in the analysis of the LGF50 gene set, although minimum gene changes in pines were significantly lower than in the LGF gene set (Fig. S3B; Table S5).

All the examined models pointed to higher rates of gene duplication and loss in Pinaceae or, whenever independently estimated, in pines, compared to angiosperms and *P. patens*. The six-λ models with highest likelihood for the LGF and LGF50 gene sets are shown in figure 4.

### Pinaceae gene turnover rates are robust to incomplete gene annotation

To verify the level of completeness in the gene sets of all analyzed species we examined protein models from each taxon with a dataset of 1,440 one-to-one orthologs found in embryophytes using the program BUSCO (Simao, et al. 2015). Flowering plants completeness ranged from ∼91.5% in banana to 99.7% in *A. thaliana*, whereas only between 52-57% of conserved proteins were encoded in the gene sets of the four Pinaceae species (Fig. 5; Table S6). The *P. patens* gene set showed an intermediate completeness level of ∼72%. Therefore, it is reasonable to assume that a large number of genes have not been identified in the four Pinaceae assemblies. This deficiency is expected to cause a significant increase in the number of gene losses along the Pinaceae lineages and to consequently inflate estimates of gene turnover rates in these species.

**Figure 5.**
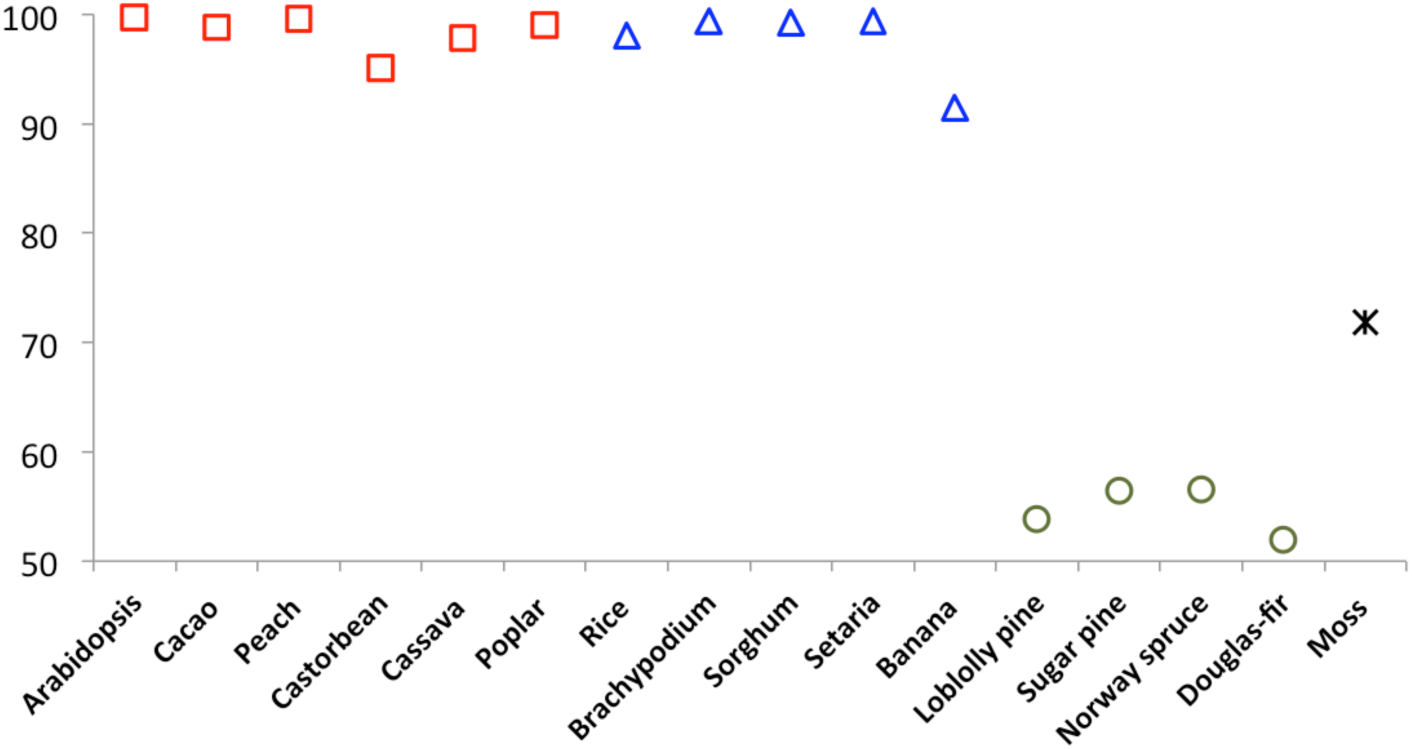
Proportion of conserved one-to-one embryophyte orthologs identified in the 16 analyzed land plants. Squares, triangles, circles and asterisk represent dicots, monocots, Pinaceae and *P. patens*, respectively.

To assess the possible effect of these “missing genes” on the observed high rates of gene turnover in Pinaceae we separately estimated gene duplication and gene loss rates in the best four-λ models from previous runs (Table S2). As expected, apparent gene losses occurred at a much higher frequency in Pinaceae compared to any other lineage (Fig. 6; Table S2). Nevertheless, Pinaceae gene duplication rates were comparable to those found in angiosperms in both the LGF gene set (Fig. 6; Table S2) and the LGF50 gene set (Fig. S4; Table S2).

**Figure 6.**
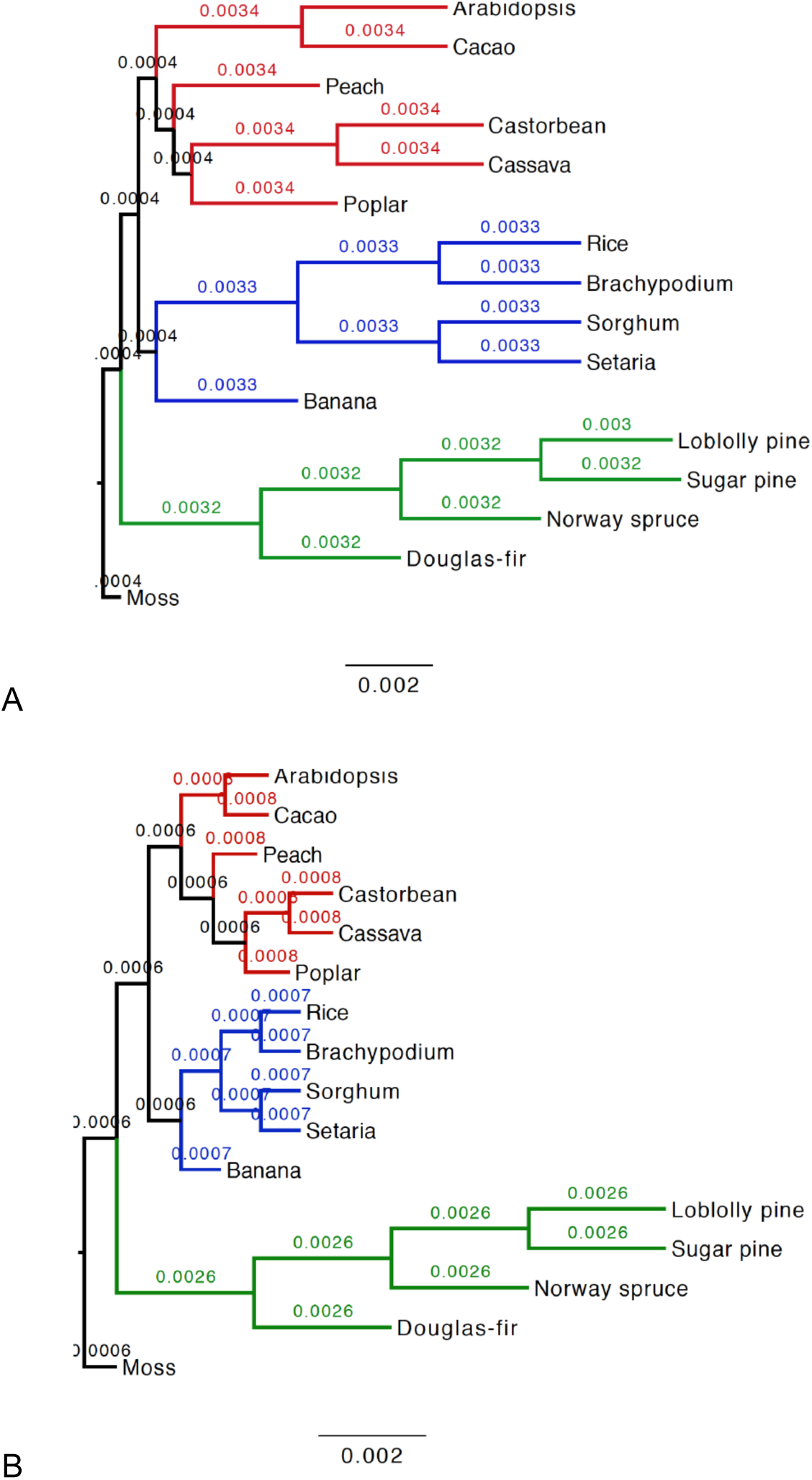
Gene duplication (A) and loss (B) rates in four-λ models applied to the LGF gene sets. Dicots, monocots and Pinaceae branches are color-coded as in Figure 1.

The six-λ models with the highest likelihood values were also analyzed (Table S3). Most models did not achieve convergence, possibly because of contrasting patterns in rates of gene gain and gene loss in some species. However, separate estimates of gene duplication and loss were obtained in at least one model for both LGF and LGF50 gene sets (Table S3; Figs. S5-S6). Notably, these results confirm an accelerated rate of both gene duplication and gene loss in pines compared to other Pinaceae, and to the combined dicot branches and monocot branches.

### Gene turnover rate estimates using Pinaceae-only gene sets

To further assess the variation in gene turnover rates among Pinaceae we determined gene gains and gene losses in the PGF and PGF50 gene sets, which included only gene families from the two pines species, the Norway spruce and the Douglas-fir (Table S1). We first analyzed all the 51 combinations of models with three possible estimates of λ values for both gene sets (Table S7). The majority of models applied to the two gene sets (97/102) reached convergence, with the highest gene turnover rates found in one or both pines (Table S7). Average gene turnover rates and gene gain/loss events were also higher in pines (Fig. 7; Table S8). Gene duplications and losses were especially elevated in sugar pine in the PGF gene set but they markedly decreased in the PGF50 set, suggesting high levels of erroneous pseudogene annotation in this species (Table S8). A minimum of 75 gene turnover events per gene per million year were estimated in the two pine species in the PGF50 gene set (Table S8).

**Figure 7.**
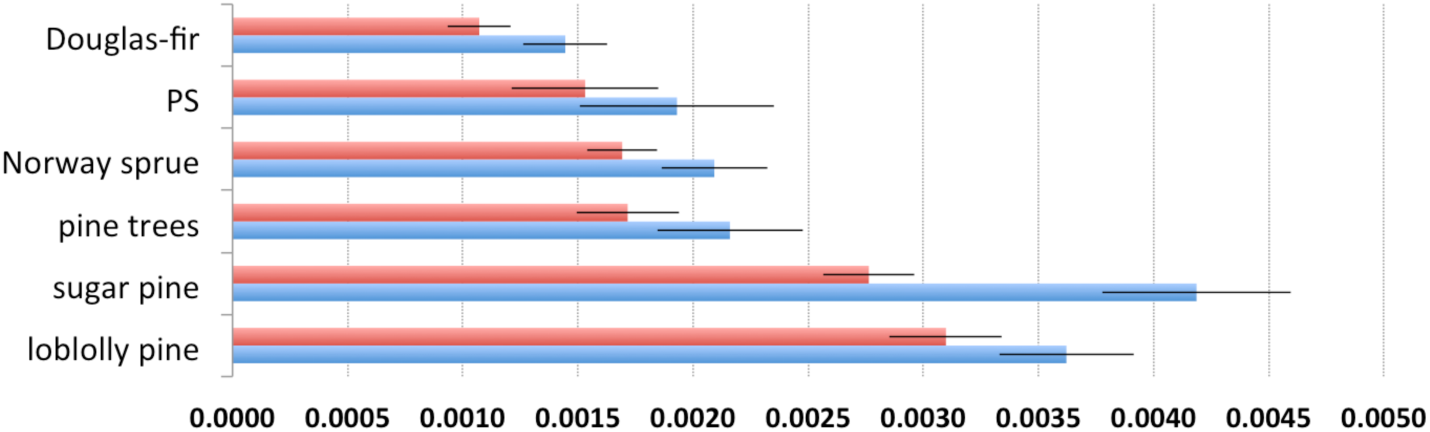
Average λ values for each Pinaceae branch in 97 three-λ models using the PGF (blue, 49 models) and PGF50 (red, 48 models) gene sets. Bars indicate 95% confidence interval. PS – pines and spruce clade.

Both PGF and PGF50 gene sets showed the highest likelihood for a model incorporating three gene turnover rate estimates for pines, Norway spruce/Douglas-fir, and the two inner branches of the phylogeny, respectively (Model M30 in Table S7). According to the estimates obtained with this model, gene turnover rates were 3.6-3.8 times higher in pines than in Norway spruce and Douglas-fir (Fig. S7; Table S7).

CAFE analyses inferring separate estimates of gene duplication and loss did not reach convergence for this model nor for other models with identical estimates for the two pine species, suggesting significant differences in the patterns of gene gains and losses between loblolly pine and sugar pine. Indeed, when separate estimates of gene duplication and loss rates were obtained for the two pine species (model M6 in Table S7), the sugar pine estimates were 1.16 to 1.41 times higher than those of loblolly pine (Figs. S8-9).

We noticed that the proportion of gene families showing opposite trends in the two pine species, that is families that expanded in one species and contracted in the other, was only ∼12% of all families with change in size in these taxa (data not shown). This suggests that a possible bias in pine gene turnover rates due to the topology of the tree—namely, the presence of two pine species vs. Norway spruce and Douglas-fir—is unlikely to affect these results. Nevertheless, we tested if the presence of two pine species was responsible of the observed gene turnover rates in the PGF and PGF50 gene sets by analyzing phylogenies with only one of the two pines (Table S9). In both cases we applied the seven possible models to the 3-species phylogenies allowing any combination of the two λ values (Table S9; Fig. S10). On average, gene turnover rates were higher in either pine species than in the other lineages (Table S10). Additionally, for both loblolly pine-only and sugar pine-only datasets, the model incorporating a separate λ value for the pine species showed the highest likelihood value, in both PGF and PGF50 gene sets (Table S9; Fig. S11). Independently estimated gene duplication rates and gene loss rates were also higher in either pine species than the rest of the phylogeny (Table S9). Taken together, these results suggest that the observed accelerated gene family evolution in pines in not due to the presence of multiple pine species in our phylogeny.

## Discussion

Differences in the protein-coding gene content between species have long been recognized as important sources of phenotypic variation. Comparative genomic studies have shown that gene duplication and gene loss events that become fixed along a given lineage are the primary cause of the observed interspecific variation in gene content in eukaryotes (Lynch and Conery 2000; Hahn, Han, et al. 2007; Carbone, et al. 2014; Soltis, et al. 2015). Species with a high frequency of gene turnover, i.e. the sum of gains and losses of genes, may enjoy an increased potential to adapt and evolve novel traits (Ohno 1970; Albalat and Canestro 2016). Therefore, determining the pace at which genes are gained and lost during species evolution, particularly in a comparative framework, is a major objective of evolutionary genomics. In their seminal works, Lynch and Conery applied steady-state demographic techniques to datasets of paralogous substitution distances (*dS* values) and found rates of gene duplication per gene per million years of ∼0.009 in humans, ∼0.001 in *Drosophila melanogaster*, ∼0.004 in *Saccharomyces cerevisiae* and *Schizosaccharomyces pombe*, ∼0.016 in *Caenorhabditis elegans*, ∼0.002 in *A. thaliana* and 0.030 in *Encephalitozoon cuniculi* (Lynch and Conery 2000, 2003). As they noted, at the time such estimates were comparable to nucleotide substitution rates at silent sites. Similar rates of gene duplication based on *dS* estimates were obtained in subsequent studies in fruit fly, budding yeast and *C. elegans* (Gu, et al. 2002), human and mouse (Pan and Zhang 2007), and *A. thaliana* (Maere, et al. 2005). On the contrary, work by Gao and Innan based on phylogenetic and syntenic approaches investigated paralogous pairs in *S. cerevisiae* and reported a much lower gene duplication rate of 0.01-0.06 per gene per *billion* years (Gao and Innan 2004). They proposed that high levels of nonallelic gene conversion had dramatically shifted the actual time of gene duplication events in budding yeast. However, other authors have pointed out flaws in this analysis, showing that gene conversion occurs at disproportionally high levels in the relatively small set of genes analyzed by Gao and Innan, thus supporting previous findings pointing to high rates of gene turnover (Lin, et al. 2006; Evangelisti and Conant 2010; Casola, et al. 2012).

Another approach to obtain genome-wide rates of gene duplication—one also used in this study—is based on modeling evolutionary dynamics of gene families along a phylogeny. Most of the implementations of this method, including CAFE, rely on maximum-likelihood inferences of gene family evolution, and provide direct estimates of gene loss (Librado, et al. 2012; Han, et al. 2013). Remarkably, estimates of gene turnover rates based on these approaches largely parallel the results generated by analyses of *dS* value distributions. For instance, the average frequency of gene turnover obtained using CAFE in budding yeasts, *Drosophila* species and mammals ranged between 0.00059 and 0.00238 per gene per million years, depending on the taxon and implementation of models including error rate estimates (Han, et al. 2013). Rates of gene gain and loss are likely higher in flowering plant lineages that experienced one or multiple polyploidization events (Tuskan, et al. 2006; Schmutz, et al. 2010; Carretero-Paulet, et al. 2015; Yang, et al. 2016). Given the current availability of genome sequences from non-model taxa, a broader view of gene turnover rates across eukaryotes is nowadays achievable. However, variation in the completeness and accuracy of gene sets remains one of main limiting factors when utilizing non-model organisms to investigate on the pace of gene gains and losses.

In this work, we developed and implemented strategies that address some of the pitfalls associated with using incomplete genome assemblies with gene sets that are still undergoing refinement to estimate gene turnover rates. We applied the maximum-likelihood framework implemented in the CAFE package (Han, et al. 2013) to analyze patterns of gene gain and loss in eleven angiosperms, four gymnosperms belonging to the Pinaceae family, and the moss *P. patens*. The Pinaceae gene sets lack up to thousands functional genes and may contain a high number of erroneously annotated short pseudogenes (Nystedt, et al. 2013; Wegrzyn, et al. 2014), thus representing an ideal taxonomic group to estimate gene duplication and gene loss events in a phylogeny in the presence of incomplete and potentially inaccurate gene annotations. We addressed these issues by filtering out short genes in Pinaceae and by separately estimating gene gain and loss rates. Additionally, we implemented models wherein multiple estimates of gene turnover rates were generated on different branches of the sixteen-plant phylogeny.

Results obtained using all genes (LGF gene set) suggested that rates of gene turnover are comparable in Pinaceae, dicots and monocots (Figs. 2-3). This finding was robust to possible occurrence of pseudogenes, as inferred using the LGF50 gene set. A bias toward gene losses was observed in Pinaceae, as expected given their gene set incompleteness. However, we noticed that gene duplication rates were similar between Pinaceae and flowering plants, particularly those with no recent WGDs, indicating that the overall high rates of gene turnover in these gymnosperms are not solely driven by an excess of apparent gene losses.

Using evolutionary models that incorporate six separate estimates of gene turnover rates along the phylogeny we were able to identify lineages within major taxonomic groups that experienced higher frequencies of gene gains and losses. Because our results were based on a phylogenetic approach, gene gains and losses due to polyploidization (whole-genome duplication) events could not be directly estimated. However, similarly to Pinaceae, cacao, castor bean and peach have not experienced recent WGDs (Van de Peer, et al. 2017). Therefore, these three “recent WGD-free” lineages offer an evolutionary parallel to the branches leading to the four Pinaceae species, which share only an ancient WGD event (Li, et al. 2015).

We sought to obtain separate rates of gene turnover in dicot lineages with and without recent WGD events in several analyzed evolutionary models (Fig. 4; Table S3). Depending on the gene set analyzed, pines showed either the highest levels of gene turnover (both minimum and maximum rates across all models) or levels that are comparable to dicots that did not experienced a recent WGD. In spite of the stringency of our filtering, particularly when generating the LGF50 gene set, we observed that gene gains and gene losses occur at a similar pace along the branches leading to pines and “recent WGD-free” angiosperm branches (Fig. 4; Table S2).

The analysis of Pinaceae-only gene sets confirmed an accelerated rate of gene turnover in pines compared to Norway spruce and Douglas-fir, with estimated frequencies of gene gains and losses ranging between 0.0028-0.0042 in loblolly pine and sugar pine (Fig. 7; Table S8). Overall, these results are in contrast to the lower nucleotide substitution rates found in Pinaceae and other gymnosperms compared to angiosperms (Willyard, et al. 2007; Buschiazzo, et al. 2012; Chen, et al. 2012; De La Torre, et al. 2017). For instance, a recent study based on 42 single-copy genes shared by seed plants indicated, on average, lower rates of both synonymous and nonsynonymous substitutions in gymnosperms compared to both woody and herbaceous angiosperms (De La Torre, et al. 2017). These authors found that average *dS*, absolute silent-site divergence (µ) and nonsynonymous substitution per nonsynonymous site (*dN*) were lower in gymnosperm vs. angiosperm families. Gymnosperms also showed markedly elevated *dN/dS* values compared to flowering plants, suggesting that adaptive substitutions are more readily fixed in the former, as previously reported in other population genetic and comparative genomic studies (Eckert, et al. 2010; Buschiazzo, et al. 2012; Eckert, et al. 2013; De La Torre, et al. 2015; Hodgins, et al. 2016)

The rate of synonymous substitution depends mainly on the rate of mutations, which form primarily during DNA replication (cell division), although mutations due to environmental factors may be more common in some plant tissues (Sarkar, et al. 2017). Because the germline is not separated from the soma in the seed plants’ sporophytic generation, some of the mutations that accumulate during mitotic cell divisions in the apical meristem will be incorporated into the gametes. A decrease in the number of meristem cell divisions in long-lived plants has been proposed to explain the observation that taller plants exhibit lower mutation rates (Lanfear, et al. 2013; Bromham, et al. 2015; Burian, et al. 2016). Furthermore, older *A. thaliana* plants have been shown to accumulate fewer mutation per unit of time than younger ones, suggesting that cells in the apical meristem that will eventually differentiate into the gametophyte may arrange into a ‘germline-like’ niche with low numbers of mitotic divisions (Watson, et al. 2016). Such properties, if shared across seed plants, could explain the low substitution rates in Pinaceae (De La Torre, et al. 2017).

Mutational mechanisms that produce gene duplication and gene loss events are associated with recombination-dependent processes (Zhang 2003), gene retrotransposition (Casola and Betran 2017), fork stalling and template switching (Zhang, et al. 2009) and changes in the chromosome number including whole-genome duplications (Conant, et al. 2014; Van de Peer, et al. 2017). These mechanisms largely differ from those generating single base pair mutations, although gene loss can also be generated through single nucleotide loss-of-funtion mutations. However, both single base pair mutations and large DNA duplications/deletions often occur during or after DNA replication and are proportional to the number of cell divisions (Arnheim and Calabrese 2009; Hastings, et al. 2009; Gaut, et al. 2011; Lu, et al. 2012). Gene retroposition may also be coupled with cell divison (Abyzov, et al. 2013). Therefore, one would expect parallel trends in the rates of single base pair mutation and DNA duplication and loss. Nevertheless, some studies have shown that these two rates may be uncoupled. For instance, hominoids (great apes) experienced both a slowdown in substitution rates due to lower frequency of mutations (Goodman 1985) and an increase in segmental duplications, particularly along the human-chimpanzee lineages (Marques-Bonet, et al. 2009).

Our results suggest a similar scenario in Pinaceae, wherein a high level of gene turnover is in contrast with low rates of nucleotide substitution. We argue that these opposite trends may be explained by elevated frequencies of gene copy number variant (CNV) formation, selection for gene family expansion and/or contraction, or a combination of these processes. Several lines of evidence, including gene birth-death estimates and mutation accumulation experiments, suggest that gene CNVs occur at rates that are a few orders of magnitude higher than per-base mutation rates in a range of prokaryotes and eukaryotes (Katju and Bergthorsson 2013). Although rates of CNV formation are not yet estimated in Pinaceae, copy-number variants seem to be common according to genome-wide studies in pines and spruces (Neves, et al. 2014; Prunier, et al. 2017). High levels of CNV formation may be fueled by illegitimate recombination between the abundant repetitive sequences, especially retroelements, in the large genomes of Pinaceae. Retroelements may also have mediated gene duplication throughout reverse transcription of mRNAs, similarly to what occurred during primate evolution (Marques, et al. 2005). Pinaceae and other conifers harbor a variety of retroelements, including some types or retroelements that are not found in other plants (Lin, et al. 2016). The analysis of the evolutionary history of genes and pseudogenes in pine, Norway spruce and Douglas-fir genomes will inform on the role of these mechanisms in the emergence of new genes in Pinaceae.

Selection has likely played a significant role in the expansion and contraction of gene families in Pinaceae as well. The elevated proportion of nonsynonymous substitutions found in Pinaceae coding regions suggest that this group has been evolving under strong selective pressures, possibly because of the efficacy of natural selection in species with large effective population sizes (De La Torre, et al. 2017). However, given the low nucleotide substitution rate in gymnosperms, the total number of nonsynonymous substitutions in Pinaceae remains relatively low compared to angiosperms. Therefore, frequent gene turnover events could have provided the primary source of novel genotypes and fostered adaptation in Pinaceae.

Finally, gene gains and losses may have boosted species diversity in Pinaceae, particularly in pines. Since its origin around 120 million years ago, the genus *Pinus* has diverged into multiple lineages, nowadays represented by ∼115 species (Willyard, et al. 2007; Saladin, et al. 2017). Postmating reproductive barriers can be established when species or populations contain alternative copies (paralogs) of pairs of recent gene duplicates, according to the divergence resolution model (Lynch and Conery 2000; Lynch and Force 2000). Thus, higher rates of gene duplication and loss in pines could have promoted the increased diversity in the genus *Pinus* compared to other Pinaceae. The possible association between species diversity and divergence resolution of gene duplicates is especially intriguing in pines given that other types of variation associated with speciation, namely karyotype changes and chromosomal rearrangements, appear to be rare in Pinaceae. For instance, all the ∼200 Pinaceae species except Douglas-fir and *Pseudolarix amabilis* share a chromosome number of 2n = 24 (Nkongolo and Mehes-Smith 2012), and a limited number of chromosomal rearrangements have been reported in this family of conifers (Pavy, et al. 2012). Comparative analyses of gene sets from a broader sample of pine species combined with estimates of gene duplication and loss timing along the *Pinus* phylogeny are warranted to better inform on the relationship between gene turnover and speciation in this diverse gymnosperm group.

## Methods

Protein sequences of ten flowering plants and the moss *P. patens* were obtained from Phytozome version 10 (https://phytozome.jgi.doe.gov/pz/portal.html). The banana dataset was retrieved from Phytozome v11. The latest available pine and Douglas-fir sets of protein sequences were downloaded from the TreeGenes genome portal in April 2016 (http://dendrome.ucdavis.edu/ftp/Genome_Data/genome/).

OrthoFinder v0.4.0 (Emms and Kelly 2015) was used to identify orthologous protein coding genes in the sixteen land plant dataset and in the four Pinaceae dataset (see section *Gene family datasets*). Orthogroups (i.e., gene families) in OrthoFinder are defined as homologous genes descendant from a single gene from the last common ancestor of the species examined. It is assumed that a parental gene of each orthogroup was present in the common ancestor of the four Pinaceae species. This method accounts for gene length and phylogenetic distance between species. The algorithm is also robust to missing genes, a potential challenge in incomplete genome assemblies. Pre-computed NCBI blastp v2.2.29+ (Camacho, et al. 2009) results were used as input for OrthoFinder. Additionally, the program mcl v14-137, an implementation of the Markov Cluster Algorithm (Enright, et al. 2002) was used by OrthoFinder with the default inflation parameter of 1.5.

The 25,446 orthogroup set obtained from the sixteen-species dataset was processed to remove gene families present in only a few species or those that showed high variation in size between species, in order to maximize the probability of achieving convergence in the maximum-likelihood analyses performed in CAFE. We retained 8,698 gene families that occurred in no less than 10 species and that showed a size standard deviation lower than 5.

In the Pinaceae-only dataset, a total of 25,002 orthogroups were identified using OrthoFinder. A total of 16,118 gene families were retained after removing families absent in Douglas-fir and families with no genes in pines and Norway spruce.

The program CAFE v3.0 was downloaded from github (https://github.com/hahnlab/CAFE). CAFE analyses on the phylogeny with one pine species were performed on gene families that occurred in at least two of the three Pinaceae species present in the tree. A total of 499 and 643 gene families had no genes in one of the pines and in Norway spruce and were thus removed from the analysis of PGF50 dataset with only one pine species.

The CAFE v3.0 allows to estimate global and species-specific error rates using the caferror.py script. However, results obtained using this script were not included in the present study for three reasons. First, error rates are modeled only for changes corresponding to a single gene gain or gene loss per gene family, while multiple gains and losses are likely to occur particularly in Pinaceae gene sets. Second, we noticed unrealistic species-specific error rates that for some models, including estimates of no error for some Pinaceae in some models. This might be due to the shortcoming of the aforementioned single gain/loss error model in the context of a phylogeny that includes distantly related taxa with high variation in gene content. Third, results obtained after including error rates in several models were quantitatively comparable to gene turnover estimates generated without applying such error rates, with the exception of decreased gene turnover rates found in dicots when using species-specific error rates. All datasets were manipulated using perl scripts and Unix commands available upon request.

The phylogenetic tree used here was based on previous studies (Jiao, et al. 2011; Bennetzen, et al. 2012; Kritsas, et al. 2012; Wang, et al. 2014; Zanne, et al. 2014). The time of divergence between lineages was based on the estimates provided in TimeTree (Kumar, et al. 2017). Sixteen-species phylogeny (numbers represent million years): (((((Arabidopsis:85,Cacao:85):27,(Peach:111,((Castor bean:15,Cassava:15):92,Poplar:107):4):1):15,(((Rice:46,Brachypodium:46):6,(Sorghum: 26,Setaria:26):26):71,Banana:123):4):223,(((Loblolly pine:90,Sugar pine:90):70,Norway spruce:160):20,Douglas-fir:180):170):100,Moss:450)

## Acknowledgments

We would like to thank Xuan Lin for assistance with perl scripts and Matt Hahn, Gregg Thomas, Mira Han for their help with CAFE models and parameters. This material is based upon work that was supported by the National Institute of Food and Agriculture, U.S. Department of Agriculture, under award number TEX0-1-9599, the Texas A&M AgriLife Research, and the Texas A&M Forest Service.

## Figure Legends

**Supplementary Table S1**. Analyzed species and genes in the Pinaceae-only datasets. Numbers in parenthesis indicate percentages of the total number of genes.

**Supplementary Table S2**. CAFE analysis with four independent gene turnover rates in the LGF and LGF50 gene sets. Lambda estimates are color-coded.

**Supplementary Table S3**. CAFE analysis with six independent gene turnover rates in the LGF and LGF50 gene sets. Lambda estimates are color-coded.

**Supplementary Table S4**. Highest and lowest gene turnover rates and gene turnover events in all lineages from the CAFE analysis with six estimates of gene turnover rates in the LGF gene set.

**Supplementary Table S5**. Highest and lowest gene turnover rates and gene turnover events in all lineages from the CAFE analysis with six estimates of gene turnover rates in the LGF50 gene set.

**Supplementary Table S6**. Completeness of land plant gene sets determined using BUSCO. Proportion of complete calculated including complete single-copy, complete duplicated and fragmented orthologs. BUSCOs refers to conserved one-to-one orthologs in embyophytes.

**Supplementary Table S7**. Highest and lowest gene turnover rates and gene turnover events in all lineages from the CAFE analysis with six estimates of gene turnover rates in the LGF gene set. Lambda estimates are color-coded.

**Supplementary Table S8**. Average gene turnover rates and gene turnover events in all lineages from the CAFE analysis with three estimates of gene turnover rates in the PGF and PGF50 gene sets.

**Supplementary Table S9**. CAFE analysis with two independent gene turnover rates in the PGF and PGF50 gene sets with one pines species only. Lambda estimates are color-coded.

**Supplementary Table S10**. Average gene turnover rates in all lineages from the CAFE analysis with three estimates of gene turnover rates in the PGF and PGF50 gene sets with one pine species only.

**Supplementary Figure S1**. A: Length distribution of genes in the LGF gene set. B: Length distribution of genes in the LGF50 gene set.

**Supplementary Figure S2**. Proportion of genes removed from the LGF gene set after applying the 50% length cutoff.

**Supplementary Figure S3**. Maximum (diamonds) and minimum (asterisks) gene turnover rates in fourteen six-λ models applied to the LGF (A) and LGF50 (B) gene sets.

**Supplementary Figure S4**. Gene duplication (A) and loss (B) rates in four-λ models applied to the LGF50 gene sets. Dicots, monocots and Pinaceae branches are color-coded as in Figure 1.

**Supplementary Figure S5**. Gene duplication (A) and loss (B) rates in six-λ models applied to the LGF gene sets. Branches are color-coded to highlight lineages with independent models imposed.

**Supplementary Figure S6**. Gene duplication (A) and loss (B) rates in six-λ models applied to the LGF50 gene sets. Branches are color-coded to highlight lineages with independent models imposed.

**Supplementary Figure S7**. Gene turnover rates in three-λ models applied to the PGF (A) and PGF50 (B) gene sets.

**Supplementary Figure S8**. Gene duplication (A) and gene loss (B) rates in the M6 model applied to the PGF gene set.

**Supplementary Figure S9**. Gene duplication (A) and gene loss (B) rates in the M6 model applied to the PGF50 gene set.

**Supplementary Figure S10**. The seven possible branch-λ parameter assignments in a phylogeny with only one pine species in two-λ model. P – pine; Ns – Norway spruce; Df – Douglas-fir.

**Supplementary Figure S11**. Likelihoods of CAFE models of Pinaceae datasets with either loblolly pine or sugar pine and two λ values. Bars show the likelihood of possible two-λ models for the seven three-species phylogenies. The two three-species phylogenies are shown above each graph in Newick format. PS: pines+spruce ancestral branch. A: Models with the loblolly pine, PGF gene set. B: Models with the sugar pine, PGF gene set. C: Models with the loblolly pine, PGF50 gene set. D: Models with the sugar pine, PGF50 gene set.

